# Single Cell Proteomics reveals partial uncoupling of the Glycerol-3-Phosphate shuttle to compensate for Electron Transport Chain dysfunction in Alexander Disease GFAP mutations

**DOI:** 10.64898/2026.10.05.756178

**Authors:** Charlotte AGH van Gelder, Elena Daoutsali, Yara Attour, Soufiane El Baghdadi, Paula Sobrevals Alcaraz, Robert M van Es, Can Gulersonmez, Edwin CA Stigter, Harmjan R Vos, Elly M Hol, Boudewijn MT Burgering

## Abstract

Alexander disease (AxD) is a rare genetic leukodystrophy caused by dominant mutations in the gene coding for the astrocytic intermediate filament protein Glial Fibrillary Acidic Protein (GFAP). Understanding disease mechanisms is complicated by the fact that mutant GFAP promotes further GFAP accumulation, making it difficult to distinguish mutation-specific effects from those caused by elevated GFAP levels. Here, we performed mass spectrometry-based single cell proteomics to investigate proteomic alterations associated with increasing levels of wildtype GFAP and a common AxD-causing mutant GFAP R239C. By analyzing individual cells spanning a broad range of GFAP expression levels, we separated dosage-dependent effects from mutation-specific responses. Cells expressing high levels of wildtype GFAP increasingly resembled R239C-expressing cells at the proteomic level, supporting a role for GFAP accumulation in disease pathogenesis. In contrast, mutant GFAP specifically correlated with increased mitochondrial antioxidant enzymes and remodeling of the proteasome tailored toward ATP- and ubiquitin-independent degradation of oxidized proteins. Integration with proteomics data from patient-derived cortical organoids revealed increased abundance of nuclear-encoded mitochondrial proteins and respiratory chain assembly factors without corresponding changes in mitochondrially encoded proteins, suggesting impaired electron transport chain (ETC) assembly. Consistent with this, AxD cells displayed reduced oxygen consumption, increased glycolytic activity, enhanced utilization of the cytosolic part of the glycerol-3-phosphate (G3P) shuttle, and accumulation of triacylglycerides and lipid droplets. We propose that ETC dysfunction promotes partial uncoupling of the G3P shuttle, allowing sustained regeneration of cytosolic NAD⁺ to support glycolysis while diverting G3P toward lipid synthesis and storage. By regenerating NAD⁺ through the G3P shuttle rather than lactate fermentation, AxD astrocytes may sustain glycolytic flux while preserving lactate production, an important metabolic substrate for neighboring neurons. Together, our findings identify ETC dysfunction as a central feature of AxD and suggest that metabolic adaptation through the G3P shuttle links mitochondrial dysfunction to glycolytic reprogramming and lipid accumulation. More broadly, these results provide new insight into how astrocyte dysfunction drives molecular pathways implicated in neurodegeneration.

## Introduction

Glial Fibrillary Acidic Protein (GFAP) is a type III intermediate filament protein predominantly expressed in astrocytes (Hol and Capetanaki 2017; Middeldorp and Hol 2011). Mutations in GFAP cause Alexander Disease (AxD), a rare and often fatal leukodystrophy characterized by the formation of GFAP-containing aggregates called Rosenthal Fibers (Brenner et al. 2001). Although AxD is caused by mutations in a single astrocytic gene, the molecular mechanisms linking mutant GFAP to astrocyte dysfunction, myelin loss, and neurodegeneration remain incompletely understood (Hol et al. 2025).

A growing body of evidence suggests that metabolic dysfunction is a central feature of AxD. Studies in cellular and animal models have reported abnormalities in mitochondrial morphology and distribution of mitochondria (Hernandez-Gerez et al. 2026; Viedma-Poyatos et al. 2022), impaired mitochondrial transfer between astrocytes and neurons (Gao et al. 2019), increased oxidative stress and lipid peroxidation (Viedma-Poyatos et al. 2022), reduced ATP production and a shift from lipid oxidation to lipid synthesis (Bellitto et al. 2025). Transcriptomic and proteomic analyses have further implicated pathways involved in energy metabolism, mitochondrial function, and oxidative stress responses (Bellitto et al. 2025; Berman et al. 2025; Heaven et al. 2022; Matusova et al. 2025). However, how mutant GFAP leads to these alterations and whether they represent primary pathogenic mechanisms or secondary adaptations remains largely unknown.

One factor complicating the interpretation of molecular phenotypes in AxD is the positive relationship between mutant GFAP expression and total GFAP abundance. Because AxD-associated mutations are typically heterozygous, affected astrocytes express both wildtype (WT) and mutant GFAP (Albee Messing 2019). Mutant GFAP induces cellular stress responses that further increase GFAP expression, potentially creating a positive feedback loop that amplifies disease progression. This observation forms the basis of the toxic-threshold model of AxD, which proposes that pathology develops when GFAP levels exceed a critical threshold, irrespective of the initiating cause. Indeed, mice overexpressing WT GFAP develop Rosenthal fibers and severe neurological abnormalities despite lacking a disease-causing mutation (A Messing et al. 1998). At the same time, growing evidence suggests that AxD-associated GFAP mutations exert mutation-specific gain-of-function effects that cannot be explained solely by increased protein abundance (Heaven et al. 2019). Distinguishing between dosage-dependent and gain-of-function effects therefore remains a major challenge in understanding AxD pathogenesis.

Most transcriptomic and proteomic studies of AxD have relied on bulk analyses that average signals across large cell populations. As a result, they cannot readily distinguish between molecular alterations that arise from elevated GFAP levels and those that are specifically driven by mutant GFAP. Single cell proteomics provides a unique opportunity to address this challenge by directly resolving proteomic variation at the level of individual cells and linking molecular phenotypes to GFAP abundance.

Here, we used mass spectrometry-based single cell proteomics to investigate the proteomic consequences of increasing levels of WT GFAP and a common AxD-causing GFAP R239C mutant. Using GFP intensity as a proxy for GFAP abundance, we analyzed cells spanning a broad range of expression levels, allowing us to distinguish dosage-dependent effects from mutation-specific responses. By integrating single cell proteomics with proteomic analyses of patient-derived cortical organoids, metabolomics, lipidomics, and functional metabolic measurements, we identify distinct effects of GFAP abundance and mutant GFAP gain-of-function. Our findings reveal mutation-associated remodeling of the proteasome and mitochondrial pathways, and implicate electron transport chain dysfunction as a potential driver of metabolic reprogramming in AxD. Together, these results provide new insights into how mutant GFAP alters astrocyte homeostasis and establish a framework for disentangling dosage-dependent and mutation-specific mechanisms in neurodegenerative disease.

## Material and methods

### Generation of eGFP-hGFAP R88C and R416W point mutations

eGFP-hGFAP WT (Addgene #169473) and eGFP-hGFAP R239C (Addgene #169476) plasmids were a gift from Federico Herrera (Letra-Vilela et al. 2020). R88C and R416W point mutations were introduced in the eGFP-GFAP WT plasmid by site-directed mutagenesis. Polymerase Chain Reaction (PCR)s were carried out using 50 ng of template DNA, 0.25 µM of mutagenesis primers and 200 µM of dNTPs in a total volume of 50 µl containing 0.5 µl of Q5 high fidelity DNA polymerase and the corresponding amplification buffer, with following amplification protocol: denaturation at 98 °C followed by 12 cycles at 98 °C for 15 s; 55 °C for 30 s; 72 °C for 8 min (1 min/kb, as specified by the manufacturer) and a final extension at 72 °C for 7 min.

PCR products were transformed in D5α competent cells via heat shock. Cells were allowed to thaw on ice and after adding 1 µg of DNA the samples were incubated for 30 minutes on ice. Heat shock was applied for 45 s at 42 °C and immediately after that 2-minute incubation on ice was performed. Cells were resuspended in 500 µL Lysogeny Broth and allowed to adapt at 37 °C for 1h. The bacteria were then grown on Muller-Hinton agar plate with selection marker ampicillin overnight at 37 °C to obtain single colonies. The following day the bacteria were grown in 200 mL LB medium with ampicillin [1:1000] and Maxi prep isolation was carried out (70-15010, Jet Star 2.0).

Following primers were used:

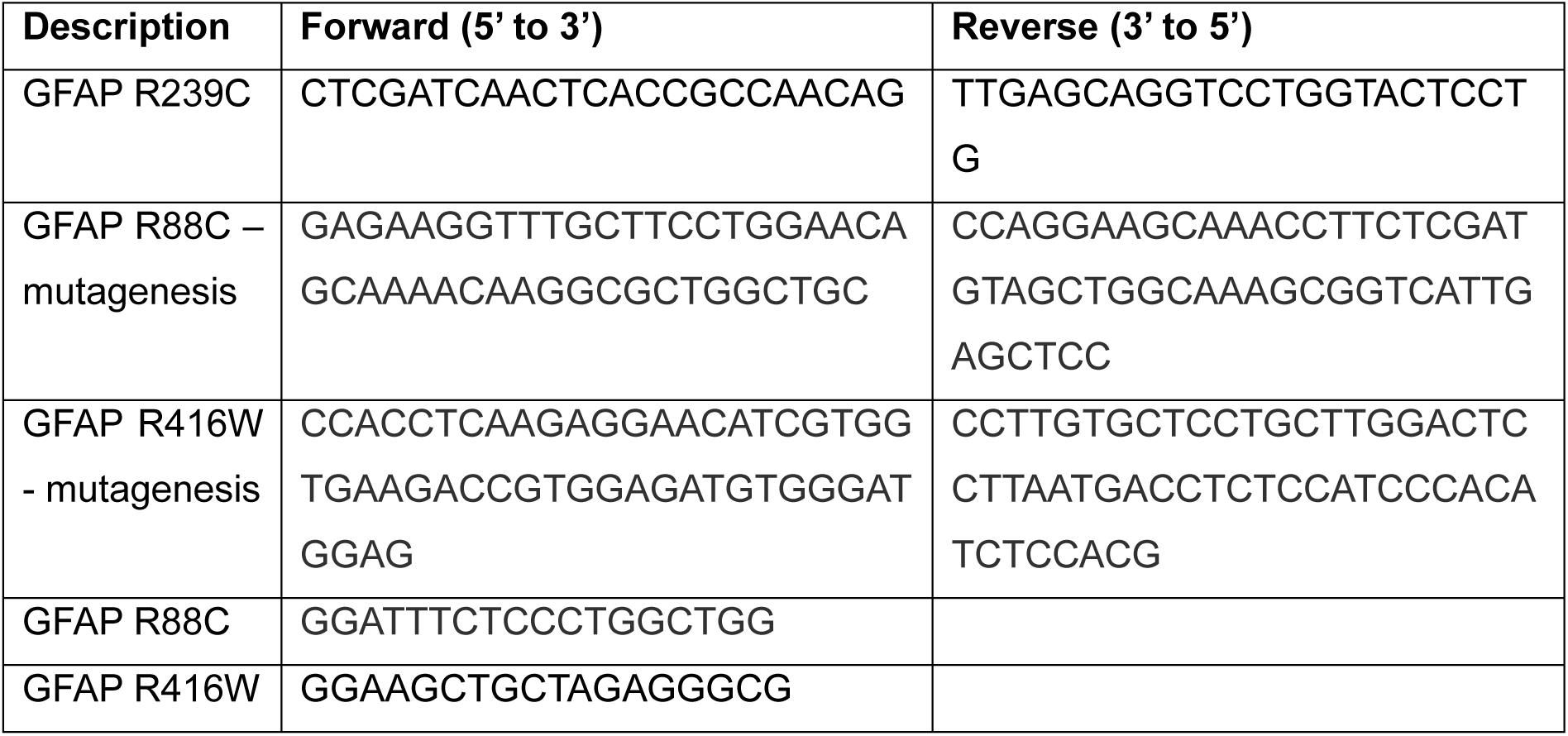

### Generation of U251-MG and U87-MG eGFP-hGFAP stable cell lines

U87-MG (ATTC accession number HTB14) and the U251-MG cell lines (high GFAP expression subclone, as in (van Bodegraven et al. 2019)) were cultured in 1:1 DMEM high glucose (D6429, Sigma-Aldrich Chemie GmbH) and F-10 nutrient mix (Gibco/Thermo Fisher Scientific, 31550-023) media supplemented with 10% Fetal Bovine Serum (FBS, S00KL10004, Bodinco BV) and 1% Penicillin/Streptomycin (P0781, Sigma Aldrich).

To create stable cell lines, U87-MG or U251-MG cells were plated in a 60 mm dish. At 70% confluency, cells were transfected with plasmids using Lipofectamine 2000 (Invitrogen) following standard protocol (8 µg cDNA and 20 µl transfection reagent). After 48 hours, selection was started by passaging the cells 1:2 and addition of 500 μg/μl G418 (Chemcruz). Monoclonal cell lines were obtained by FACS sorting of single eGFP positive cells (BD FACSAria III) in a 96 wells plate and subsequent expansion. The single cell proteomics experiments were performed on the U251-MG cell lines; all other experiments were performed with the U87-MG cell lines.

### Generation of induced Astrocytes

AxD iPSCs, derived from a 6-year-old AxD patient carrying the heterozygous point mutation in GFAP (c.715C > T, p.R239C), and isogenic CRISPR/Cas9-corrected control iPSCs were received from Natasha Snider (Department of Cell Biology and Physiology, University of North Carolina at Chapel Hill, USA) and were described previously (Battaglia et al. 2019). iPSCs were transduced and differentiated as described earlier (Canals et al. 2018). Briefly, iPSCs were transduced with 3 lentiviruses (3rd generation) containing the *reverse tetracycline-controlled SRY-Box Transcription Factor 9* (*Sox9*; Addgene, 117269), the *Nuclear Factor I B* (*Nfib*; Addgene, 117271), and the *transactivator* (*rtTA*; Addgene, 20342) respectively. iPSCs that reached 80% confluency, without signs of spontaneous differentiations, were dissociated with Accutase (StemCell Technologies, 07922) for 5-7 minutes at 37°C and single cells were plated in Geltrex (ThermoFisher Scientific A1413202)-coated 6-well plates (Corning, 354230) in StemFlex medium (ThermoFisher Scientific, A3349401) supplemented with 5 µM ROCK inhibitor (Axon Mechem, Axon 1683). One day later, iPSCs were transduced with the 3 lentiviruses (MOI ∼1.5) in StemFlex medium. On day 0, StemFlex medium was replaced with StemFlex medium supplemented with doxycycline (2.5 µg/mL; Sigma-Aldrich, D9891). On day 1 and 2, cells were cultured in Expansion medium containing; DMEM/F-12 (Thermo Fisher Scientific, 11320074), 10% Fetal Bovine Serum (FBS; Sigma-Aldrich, F7524), 1% N2 supplement (Thermo Fisher Scientific, 17502001), 1% L-glutamine (Thermo Fisher Scientific, 25030024) and supplemented with 2.5 µg/mL doxycycline, 1.25 µg/mL puromycin (ThermoFisher Scientific, A1113803), and 100 µg/mL hygromycin (ThermoFisher Scientific, 10687010). From day 3 to day 6, the Expansion medium was gradually replaced with FGF medium containing; Neurobasal medium (ThermoFisher Scientific, 21103049), 2% B27 (Thermo Fisher Scientific, 17504001), 1% MEM Non-Essential Amino Acids Solution (NEAA; Thermo Fisher Scientific, 11140035), 1% L-glutamine, and 1% FBS, and supplemented with 8 ng/ml human fibroblast growth factor basic (hbFGF; Peprotech, 100-18B), 5 ng/ml ciliary neurotrophic factor (CNTF; Peprotech, 450-13), 10 ng/ml Bone morphogenetic protein 4 (BMP4; Peprotech, 120-05ET), and 2.5 µg/mL doxycycline. Hygromycin (100 µg/mL) was added to the medium until day 5.

On day 7, iAstrocytes were dissociated with accutase and replated in FGF medium supplemented with 8 ng/ml hbFGF, 5 ng/ml CNTF, 10 ng/ml BMP4, and 5 µM ROCK inhibitor in the desired density on Growth Factor Reduced (GFR)-reduced Matrigel (CLS35430, Merck)-coated plates. On day 8, FGF medium with supplements (without ROCK inhibitor) was refreshed. From day 9 to day 21, half of the medium was replaced by Maturation medium containing; 1:1 DMEM/F-12 and Neurobasal, 1% N2, 1% sodium pyruvate (Thermo Fisher Scientific, 11360070), 1% L-glutamine, 5 µg/ml N-acetyl-cysteine (Sigma-Aldrich, A8199), and supplemented with 500 µg/ml N6,2′-O-Dibutyryladenosine 3′,5′-cyclic monophosphate sodium salt (dbcAMP; Sigma-Aldrich, D0627), 5 ng/ml heparin-binding EGF (Sigma-Aldrich, E4643), and 10 ng/ ml CNTF, 10 ng/ml BMP4. On day 21, iAstrocytes were collected for subsequent experiments.

Media for induced astrocytes at the different time points.

|  |  |
| --- | --- |
| Day 0 | SF |
| Day 1-2 | Expansion |
| Day 3 | 75% Expansion + 25% FGF |
| Day 4 | 50% Expansion + 50% FGF |
| Day 5 | 25% Expansion + 75% FGF |
| Day 6-8 | FGF medium |
| Day 9-21 | Maturation medium |

### Mass Spectrometry of single cells

#### Preparation of single cell digests

U251-MG eGFP-GFAP and eGFP-R239C-GFAP were washed with phosphate buffered saline (PBS), released from culture dish with trypsin, quenched with full growth media, and spun down at 1200 rpm for 3 min. Cell pellets were resuspended in ice-cold PBS and washed twice to remove any remaining growth medium before cell sorting. Cell sorting was performed on the Pala Cell Sorter (Bio-Techne) using a single cell microfluidic cartridge (Bio-Techne) into 96-well plates preloaded with 1 μl 2x concentrated lysis and digestion buffer consisting of 0.4% n-Dodecyl-β-D-maltoside (DDM) (Cayman Chemical Company) and 2 ng/μl LysC&Trypsin (Promega) in 200 mM triethylammonium bicarbonate buffer (TEAB) pH 8.5 (Sigma-Aldrich). Plates containing sorted cells were covered with a silicone mat (Thermo Fisher Scientific) and digested in a thermal cycler with heated lid to avoid evaporation at 50 °C for 90 min. Samples were acidified with 3 μl of 0.1% formic acid.

#### Liquid Chromatography and Mass Spectrometry Data Acquisition

Peptides were chromatographically separated on a Vanquish Neo UHPLC system connected to a 25 cm in-house made pulled emitter fused silica column 519 (50 µm ID, Polymicro) packed with 1.9 µm aquapur gold C-18 material (dr. Maisch) in trap-elution configuration. A 20-min method was utilized, where buffer B was increased from 7% to 30% (10 min), 30% to 60% (6 min) and 60% to 100% (2 min) with the nominal flow set at 400 nl/min. The flow was kept constant for 2 min, after which the run was stopped and the column equilibrated.

Data acquisitionwas conducted with an Orbitrap Astral mass spectrometer operated in positive mode with the FAIMS Pro interface (Thermo Fisher Scientific) using a compensation voltage set to −48 V. MS1 spectra were acquired with the Orbitrap at a resolution of 480,000 and a scan range of 400 to 800 m/z with normalized automatic gain control (AGC) target of 300% and maximum injection time of 60 ms. Data-independent acquisition of MS2 spectra was performed in the Astral using loop control set to 1.2 s per cycle with an isolation window of 10 m/z and maximum injection time of 50 ms. A 1 m/z window overlap and a range of 400 to 800 m/z was used. Fragmentation of precursor ions was performed using higher energy collisional dissociation (HCD) using a normalized collision energy (NCE) of 25% and AGC target was set to 500%.

#### Mass Spectrometry RAW Data Analysis

The single cell RAW data was searched with Biognosys Spectronaut Software [version 20.1.250624.92449], using directDIA analysis. Protein identification was performed against the UniProt *Homo sapiens* reference proteome fasta file (taxonomy ID: 9606; downloaded September 2024), common contaminants were also searched for. Trypsin/P with a maximum of two missed cleavages was set as digestion enzyme and acetyl (protein n-term) and oxidation (m) were set as variable modifications. The BGS Factory Settings were used for the analysis, with a q-value cutoff set at 0.01 on both protein and peptide level. Protein quantification was set at MS1, with the LFQ algorithm set for MaxLFQ. Cross-Run Normalization was selected using the automatic normalization strategy.

#### Data Filtering and Analysis

The protein quantification matrix report was exported, and the output file was further analyzed in R [version 4.3.3]. Cells with a low amount of protein identifications were excluded using a cut-off based on the number of proteins found in controlled empty runs. Potential contaminant proteins were filtered out and to ensure robust quantification, proteins with excessive missing values were removed by allowing an identification threshold of maximum 50% missing values across all cells, missing values were then imputed for further downstream analysis using a manually defined left-shifted and scaled imputation based on the normalized log2 data distribution. To identify proteins showing significant differences across the GFAP intensities, analysis of variance (ANOVA) was performed for each protein using the *stats* package in R. P-values were adjusted for multiple testing using Benjamini-Hochberg algorithm to control for false discovery rate (FDR). Proteins with an adjusted p-value smaller than 0.05 were considered statistically significant, and the topmost significantly different proteins were selected for dimensionality reduction analysis. The selected significant proteins were used for both Uniform Manifold Approximation and Projection (UMAP) and Heatmap visualizations, which were performed using the *uwot and ComplexHeatmap* packages in R, respectively. These visualizations together with publicly available datasets (MitoCARTA version 3.0 (Rath et al. 2021)) allowed for the identification of cells clustering patterns, as well as characterization of specific groups of co-regulated protein expressions. To identify potential interesting correlating proteins, correlation analysis was performed. For cells in which the GFAP was detected, Spearman correlations were computed between the target protein and other protein subsets using pairwise complete observations, requiring a minimum of four valid sample pairs per test. For each protein, the correlation coefficient (ρ), the p-value and the number of observations were recorded. P-values were adjusted for multiple testing using the Benjamini-Hochberg method, and proteins were ranked based on their correlation strength. To provide a clear view of their association, several protein subsets were plotted against the target protein (GFAP) and linear regression was used to further quantify their correlations.

### Mass Spectrometry of induced astrocytes

#### Sample preparation

Astrocytes were released from culture plates with Accutase and pelleted by centrifugation for 4 min at 200 g. Pellets were washed twice with PBS and dry pellets were flash frozen and stored at −80 °C until further processing. Proteins were lysed in 0.2% DDM (Cayman Chemical Company) and 100 mM TEAB (Sigma-Aldrich) and incubated on ice for 10 min, after which lysates were spun down at 14,000 rpm for 10 min at 4 °C. Supernatants were digested with 2 ug/ul LysC&Trypsin (Promega) for 2 hours at 50 °C and acidified to a final concentration of 1% formic acid.

#### Liquid Chromatography and Mass Spectrometry Data Acquisition

Peptides were chromatographically separated on a Vanquish Neo UHPLC system connected to a 25 cm in-house made pulled emitter fused silica column 519 (50 µm ID, Polymicro) packed with 1.9 µm aquapur gold C-18 material (dr. Maisch) in trap-elution configuration. A 60-min method was utilized, where buffer B was increased from 7% to 20% (36 min), 20% to 40% (18 min) and 40% to 100% (3 min) with the nominal flow set at 400 nl/min. The flow was kept constant for 3 min, after which the run was stopped and the column equilibrated.

Data acquisitionwas conducted with an Orbitrap Astral mass spectrometer operated in positive mode with the FAIMS Pro interface (Thermo Fisher Scientific) using alternating compensation voltages of −45 and −65, respectively. MS1 spectra were acquired in DIA mode with the Orbitrap at a resolution of 240,000 and a scan range of 390 to 980 m/z with normalized automatic gain control (AGC) target of 500% and maximum injection time of 5 ms. Data-independent acquisition of MS2 spectra was performed in the Astral using loop control set to 0.6 s per cycle with an isolation window of 2 m/z and maximum injection time of 3.5 ms at a m/z range of 400 to 800. Fragmentation of precursor ions was performed using higher energy collisional dissociation (HCD) using a normalized collision energy (NCE) of 25% and AGC target was set to 500%.

### Mass spectrometry analysis of metabolites and lipids

#### Materials

Organic solvents were ULC-MS grade and purchased from Biosolve (Boom B.V., Meppel, The Netherlands). Chemicals were analytical grade and purchased from Sigma-Aldrich (Zwijndrecht, The Netherlands). Ultrapure water was obtained fresh from a Milli Q instrument (Merck Millipore, Amsterdam, The Netherlands). A Labconco Centrivap and thermo shaker were obtained from VWR International (Amsterdam, The Netherlands). A Vanguard UPLC system, an Exploris 480 Orbitrap MS, Compound Discoverer v3.3 software and XCalibur software v4.7 were purchased from Thermo Scientific (Breda, The Netherlands). UPLC columns were obtained from Waters (Etten-Leur, The Netherlands).

#### Liquid Chromatography Mass Spectrometry (LC-MS)

Sample fractions in 1.5 mL Safe-Lock cap Eppendorf tubes were evaporated to dryness in a Centrivap sample concentrator at 40°C. To the dry samples 200 µL methanol (containing 50 µM BHT) and 400 µL chloroform were added. The samples were homogenized by ultrasonification for 5 min and incubated at for 1 h at 30°C and 900 rpm in a thermoshaker. Phase separation was induced by adding 200 µL ultrapure water. The samples were pulse vortex mixed, stored for 10 min at 4°C, and centrifugated for 10 min at 4°C and 17000 x g. Both the upper aqueous phase and lower organic phase were collected in labelled Eppendorf vials and evaporated to dryness in a Centrivap concentrator and stored at −80 until analysis.

On the day of analysis the sample residues were dissolved in 100 µL milli Q water and transferred to sample vials kept at 5°C during analysis. Quality control (QC) samples were prepared by pooling 10 µL aliquots from each individual sample. QC samples were analyzed after every tenth experimental sample throughout the sample analysis sequence.

For reversed phase separation either an Atlantis Premier BEH C18 AX UPLC-column (2.1×100 mm, 1.7 μm) for analysis at basic conditions, and an HSS-T3 UPLC-column (2.1x100 mm, 1.8 µm) for analysis at acidic condition. The reversed phase columns were operated at a flow rate of 300 µL min^-1^ and kept at 30°C. Mobile phase A for analysis at basic conditions consisted of 20 mM ammonium acetate and 5 mM ammonium hydroxide, and at low pH it consisted of 0.1% formic acid. Mobile phase B was acetonitrile in both cases. A step gradient was started upon injection of 5 µL of sample. The gradient steps were 0% B for 1.5 min followed by a 5 min linear increase to 15% B after which the percentage B was increased linearly to 70% during 3.5 min. The gradient was kept at 70% for 2 min, after which the system returned to 0% B in 0.2 min and the column was allowed to re-equilibrate for 7 min prior to a next injection.

For lipidomics analysis the residue of the organic phase was dissolved in 100 µL acetonitrile. Analysis was conducted with the system described above using an Acquity BEH C18 column (2.1x100 mm, 1.7 µm) kept at 60°C. The system was operated at a flow rate of 450 μL min^−1^. The mobile phases consisted of 40% acetonitrile also containing 10 mM ammonium acetate (solvent A), and 10% acetonitrile −90% isopropanol also containing 10 mM ammonium acetate (solvent B) for both negative and positive mode. A 12-min linear gradient of 40–100% B was started after the injection of 5 µL of the sample. The system was kept at 100% B for the next 5 min, after which the system returned to its starting situation. Total runtime was 20 min.

The column outlet was coupled to the Exploris 480 MS. A heated electrospray ionization (HESI) interface was used for ionization. Data were acquired in both negative and positive ionization modes, with spray voltages of 3.6 kV and 2.5 kV, respectively. The capillary temperature was maintained at 320 °C, the auxiliary gas at 11 arbitrary units, the spare gas to 2 arbitrary units, and the S-lens RF level to 50. Spectra were acquired in full-scan mode using the Orbitrap analyzer at a resolution of 180,000, with an automatic gain control of 3 x 106, a maximum injection time of 200 ms, and a mass accuracy of 5 ppm. MS/MS acquisition was performed using a collision energy of 40. For metabolite identification, a QC and standards sample were additionally analyzed by MS/MS. Data analysis was performed using XCalibur Quan and Compound Discoverer 3.3 software (Thermo Scientific), using data from standards and the KEGG, HMDB and lipidmaps databases.

### Seahorse XF Metabolic Flux Analyses

The Seahorse XF Analyzer (Agilent) was used for the Mito Stress Test and Glycolysis Stress Tests. One day before analysis, cells were seeded into XF24 V7 PS plates precoated with rat tail collagen (Corning) at 50,000 cells per well and incubated at 37 °C / 6% CO2. For the Mito Stress Test, cells were incubated in Seahorse XF medium supplemented with 10 mM glucose, 1 mM Na-pyruvate, and 2 mM L-glutamine one hour prior to start of the experiment, unless specified otherwise. Oxygen Consumption Rate (OCR) and Extracellular Acidification Rate (ECAR) were monitored upon serial injections of oligomycin (1 µM), FCCP (1 µM), and Rotenone and Antimycin A (1 µM). For the Glycolysis Stress Test, cells were incubated in Seahorse XF medium supplemented with 2 mM L-glutamine one hour prior to start of the experiment. OCR and ECAR were measured upon serial injections of glucose (10 mM), oligomycin (1.5 µM) and 2-Deoxy-D-glucose (2-DG, 50 mM). OCR and ECAR values were normalized to the final protein amount in each well, as determined by Pierce BCA protein assay (Thermo Scientific).

### siRNA experiments

Cells were transfected with ON-TARGETplus SMARTPool siRNA targeting human GPD1 (Dharmacon, Entrez Gene 2819) or scrambled sequence (Dharmacon, sequence UAGCGACUAAACACAUCAAUU) using Lipofectamine RNAiMAX Transfection reagent (Invitrogen) following manufacturer’s instructions. In short, cells were seeded at 50,000 (Seahorse) or 75,000 (other experiments) and transfected with 10 or 30 pmol siRNA in Opti-MEM medium containing 3 or 9 µl of RNAiMAX reagent right after plating. Cell culture media was refreshed four hours after transfection to avoid toxicity. All experiments were performed 24-48 hours after transfection.

### Western Blot

Protein lysates were obtained by washing cells with PBS and subsequently lysed and scraped in sample buffer (2% SDS, 5% 2-mercaptoethanol, 10% glycerol, 0.002% bromophenol blue, 300 mM Tris-HCL, pH 6.8). After denaturation at 95 °C for 5 min, 20 μg of protein was loaded onto SDS-PAGE and transferred to Immobilon PolyScreen PVDF transfer membranes. Samples were blocked in TBS-Tween (1% v/v) containing 2% Bovine Serum Albumin for 1h at 4 °C. Primary antibodies included rabbit α-GPD1 (Proteintech 13451-1-AP) and mouse α-PCNA (Invitrogen MA5-11358). These were applied in TBS-Tween 0.1% at a dilution of 1:2,000 at 4 °C overnight followed by 1h incubation of secondary HRP-conjugated antibodies (1:10,000) targeting either mouse or rabbit IgG in TBS-Tween 0.1% at RT. Protein abundance was detected by enhanced chemiluminescence method using ImageQuant LAS (GE HealthCare) with ImageQuant LAS4000 (version 1.3) software. Contrast was adjusted for clarity by linear image processing in Adobe Photoshop if required.

### Flow cytometry

Cells were plated at a density of 75,000 cells per well in a 12-well format in standard culturing conditions 24 hours prior to analysis. To measure mitochondrial superoxide production, cells were incubated with 5 μM MitoSOX (Invitrogen) in PBS for the final 30 min at 37 °C. Then, cells were released from the culture plate with trypsin, quenched with abundant FBS-containing media, and spun down at 1200 rpm for 3 min. Cell pellets were resuspended in ice cold PBS and kept on ice until analysis. Flow cytometry was performed using a BD FACS Celesta #660345. Only GFP positive cells were selected for analysis, and a minimum of 10,000 cells were analyzed per sample.

### Determination of lipid droplets

Cells were seeded into a flat bottom 96-wells plate at 6,000 cells per well, 24 hours prior to analysis. Right before analysis, cells were incubated with LipidSpot 610 (Biotum) 1:10,000 for 30 min at 37 °C, after which fluorescence was measured at 592/638 ex/em on a Spectramax M3 (Molecular Devices). Wells with cells but without LipidSpot served as a negative control.

For live cell imaging, cells were seeded at 25,000 cells per well in 8-wells μ-Slides (Ibidi) 24 hours before analysis. 30 minutes before analysis, cells were incubated with LipidSpot610 (Biotum, 1:10,000). Live cell imaging of lipid droplets was performed on a SP8 confocal microscope (Leica Microsystems) with a 40x oil-water immersion objective. Hoechst was excited at 405 nm and emission measured at 415-481 nm. LipidSpot610 was excited at 592 nm and emission measured at 638 nm. Z-stacks were acquired and displayed as maximum projections using Fiji.

## Results

### Single cell proteomics analysis of GFAP R239C cells reveals proteasomal remodeling to facilitate the degradation of oxidized proteins

Higher levels of mutant R239C GFAP correlate with earlier age of onset and more severe disease progression (Bachetti et al. 2026; Prust et al. 2011). To investigate which molecular changes are correlated with increased GFAP abundance, we performed mass spectrometry-based single cell proteomics analysis of U251-MG astrocytoma cell lines stably overexpressing eGFP-GFAP, hereafter referred to as GFAP WT, or eGFP-GFAP-R239C, respectively. We sorted almost 100 single cells per cell line into four bins with increasing GFP intensity (Figure 1A, S1A). Mass spectrometry-based quantification resulted in a dynamic range of three orders of magnitude difference between the lowest and highest GFAP abundance (Figure 1B, S1B). Uniform Manifold Approximation and Projection (UMAP) of the single cells showed clustering based on GFAP intensity (Figure 1C). Interestingly, while lower levels of WT and mutant GFAP separated in the first dimension, cells with the highest levels of WT GFAP clustered with mutant GFAP (Figure 1D). Indeed, previous studies have found that high levels of WT GFAP in mice result in similar phenotypes as GFAP mutations in models of Alexander Disease (A Messing et al. 1998).

**Figure 1.**
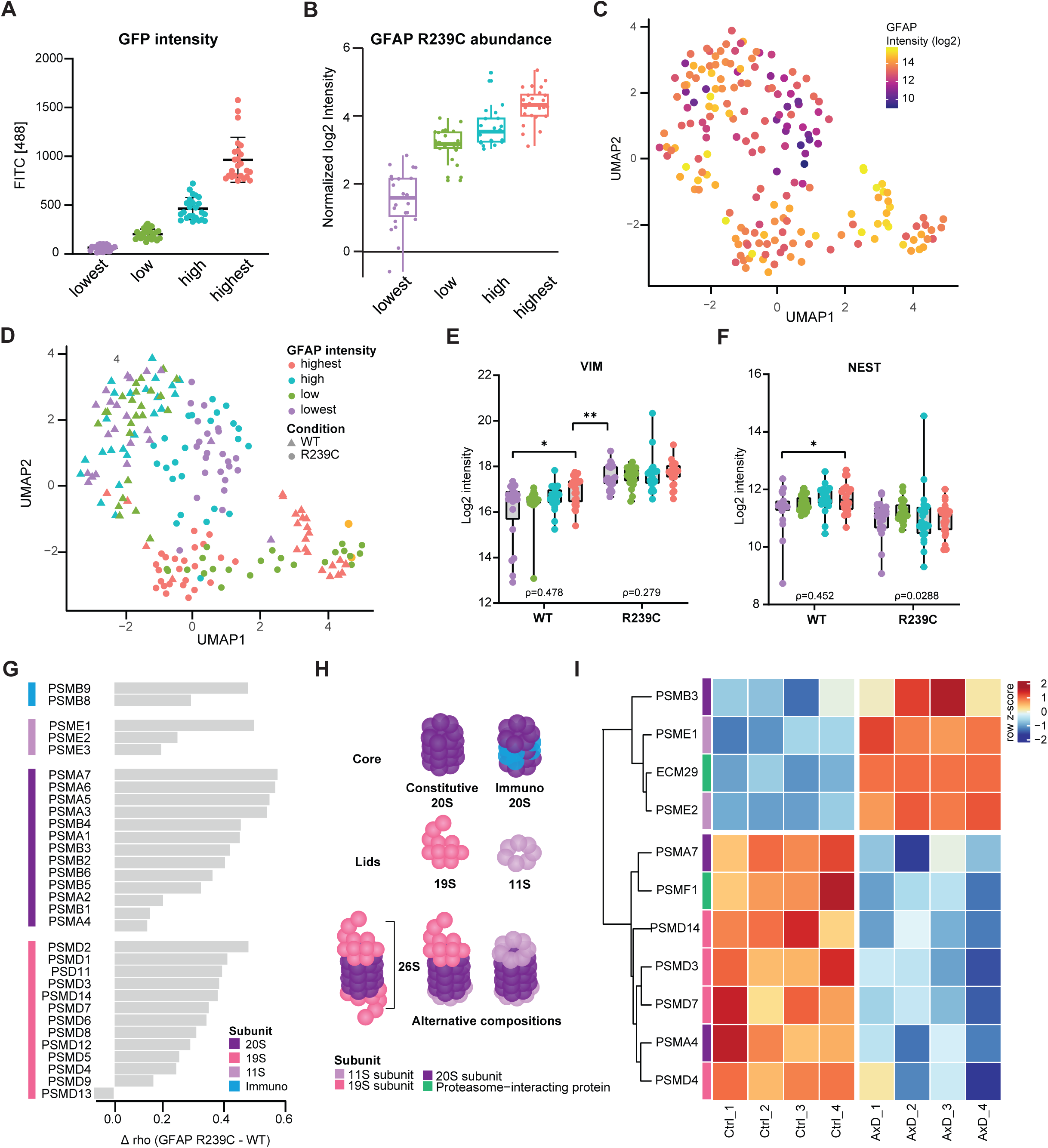
Single cell proteomics analysis of GFAP R239C cells reveals proteasomal remodeling to facilitate the degradation of oxidized proteins. **(A)** Single cells were sorted based on eGFP-GFAP-R239C into four bins (n=24 per bin). **(B)** Mass spectrometry-based quantification of GFAP protein of the single cell samples. **(C)** UMAP projection of single cells shows separation on GFAP abundance. **(D)** Wildtype GFAP cells with the highest amount of GFP intensity cluster with GFAP R239C cells. **(E)** GFAP abundance correlates well with other intermediate filament proteins Vimentin and **(F)** Nestin. The correlation is largely lost in GFAP R239C cells. T-test p-values: * p < 0.05, **p < 0.01. **(G)** Difference in correlation between GFAP WT and R239C with different proteasome subunits. **(H)** Schematic overview of different proteasome subunit compositions. Figure adapted from (Cromm and Crews 2017). **(I)** Heatmap containing proteasome subunits with significantly different abundance (t-test p-value <0.01) between AxD and isogenic control organoids.

To evaluate whether our single cell proteomics approach could identify biologically relevant protein co-regulation patterns across a range of GFAP expression levels, we correlated GFAP abundance with all other quantified proteins (Figure S1C). As a proof-of-principle, GFAP abundance in WT cells strongly correlated with vimentin (ρ = 0.452) (Figure 1E), consistent with their well-established cooperative role in the assembly of the intermediate filament scaffold (Hsiao et al. 2005). This correlation was much lower in GFAP R239C cells (ρ = 0.279), suggesting disruption of the coordinated regulation normally observed between these two intermediate filament proteins. In addition, vimentin abundance was elevated in GFAP R239C cells irrespective of GFAP levels, likely reflecting cytoskeletal remodeling associated with impaired filament assembly, a hallmark of AxD. A similar pattern was observed for the intermediate filament protein nestin, for which the positive correlation with GFAP was diminished in GFAP R239C cells despite an overall reduction in nestin abundance compared with WT cells (Figure 1F). Together, these findings demonstrate that our single-cell correlation strategy captures known biological relationships and is sufficiently sensitive to detect disease-associated alterations in protein co-regulation.

We then sought to identify proteins that gained a positive correlation in GFAP R239C cells compared to WT, by calculating the difference in correlation (delta rho). Many subunits of the proteasome complex were correlated positively with R239C GFAP abundance, including subunits of the immunoproteasome and 11S subunits (Figure 1G). The 26S proteasome, comprised of the 20S core particle flanked with two 19S caps, is the predominant form of the proteasome found in eukaryotic cells, and is responsible for ATP-dependent breakdown of ubiquitinated proteins (Cromm and Crews 2017; Thibaudeau and Smith 2019). However, the 20S particle can function in different compositions, depending on cellular demand (Figure 1H). We identified specific co-regulation of mutant GFAP with several immunoproteasome subunits, as well as subunits of the 11S lid, both of which are known to be upregulated during cellular stress (Cromm and Crews 2017).

We compared these data to our previously published bulk proteomics experiments from 150-day old cortical organoids generated from induced pluripotent stem cells (iPSCs) from an AxD patient carrying the GFAP R239C mutation (Matusova et al. 2025). AxD organoids showed a significant decrease in 19S subunits compared to isogenic controls, and an increase in 11S subunits (Figure 1I). Moreover, there was an increase in ECM29, an adapter protein that has been shown to disassemble the 26S proteasome in response to oxidative stress (Wang et al. 2017). Together, these data indicate proteasomal remodeling to facilitate the degradation of oxidized and damaged proteins.

### The AxD R239C mutation results in disbalance between mitochondrial- and nuclear encoded mitochondrial proteins, affecting electron transport chain assembly and function

We next focused on proteins that correlated most with GFAP R239C abundance. These included proteins involved in mitochondrial redox balance (CH60, LONM, PRDX3) (Figure S2A), that were significantly less abundant in GFAP R239C cells but increased with increasing GFAP R239C levels. Interestingly, we saw an increase in abundance in subunits of the electron transport chain (ETC) complexes with increasing GFAP R239C abundance. More specifically, all subunits of the F(1) sub-complex of complex V showed increased correlation with GFAP abundance (Figure 2A). This complex is responsible for the generation of ATP.

**Figure 2.**
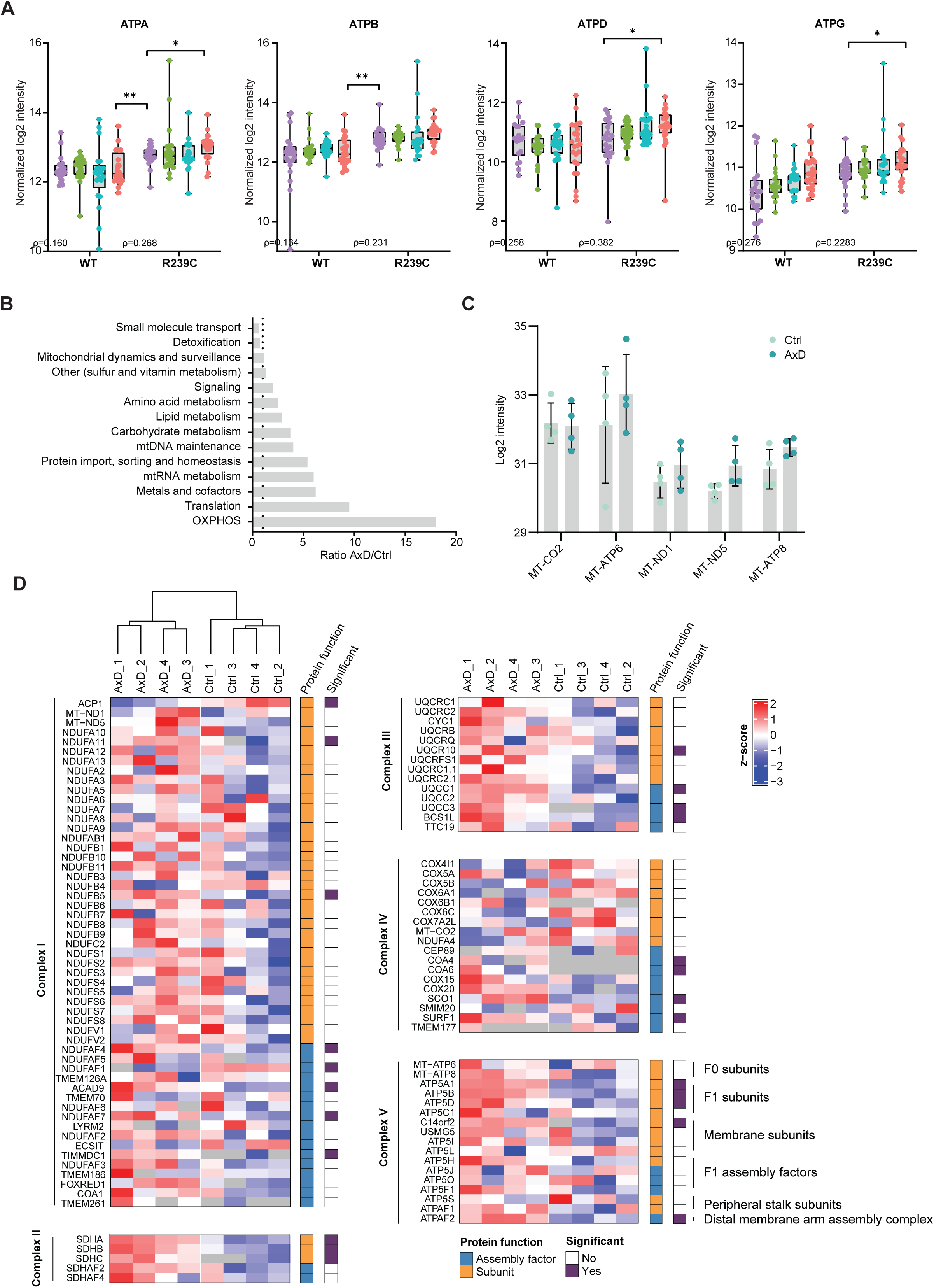
The AxD R239C mutation results in disbalance between mitochondrial- and nuclear-encoded mitochondrial proteins, affecting electron transport chain assembly factors. **(A)** The GFAP R239C mutation results in increased correlation of GFAP with subunits of complex V of the ETC. **(B)** AxD cortical organoids contain an increased number of proteins with higher abundance of mitochondrial proteins involved in oxidative phosphorylation, metabolism and mitochondrial homeostasis, while proteins involved in detoxification and small molecule transport were decreased compared to isogenic controls. **(C)** Relative abundance of detected mitochondrially encoded proteins in AxD and isogenic control organoids. **(D)** Heatmap containing all identified proteins of the electron transport chain protein complexes in AxD and isogenic control cortical organoids (t-test p-value <0.01). Proteins were divided into complex subunits (orange) and assembly factors (blue).

Similarly, the AxD organoids proteomics data showed a marked increase in abundance of many nuclear-encoded mitochondrial proteins. RNA transcripts of these proteins showed a similar trend, although these differences were not statistically significant, indicating that these differences were not originating from transcriptional origin (Figure S2B). Moreover, no differences were observed between mitochondrial and cytosolic ribosomal proteins, indicating that there is no disbalance in translation (data not shown).

We used the MitoCARTA database of mitochondrial proteins to investigate in what processes these proteins were involved and found that the proteins with increased abundance in AxD organoids were mostly related to oxidative phosphorylation, the mitochondrial central dogma, metals and cofactors, and protein import and homeostasis (Figure 2B). We did however not observe significant differences in abundance of mitochondrially encoded (MT) proteins we identified (Figure 2C), nor in proteins involved in mitochondrial fission and fusion (Figure S2C). As all MT proteins are essential components of the ETC, we performed hierarchical clustering of all proteins involved in oxidative phosphorylation which included both subunits and assembly factors of all five complexes of the respiratory chain. AxD organoids showed a significant increase in most assembly factors of complexes I, III, IV and V, but not of complex II (Figure 2D). Interestingly, complex II is the only ETC complex that does not contain MT subunits. In contrast, all three subunits of complex II were upregulated in AxD organoids, which might reflect their dual role as both ETC and TCA cycle enzymes. Complex V subunits of the F(1), or ATPase, were significantly increased while the F(0), or proton pump complex, was not affected.

As assembly factors are involved in the formation of (sub)complexes that contain both MT and nuclear-encoded proteins, we hypothesized that the increased abundance of these assembly factors might reflect a defect in ETC assembly caused by a disbalance in MT and nuclear encoded proteins, leading to ETC deficiency. To investigate this, we generated stable cell lines overexpressing two additional AxD-causing GFAP mutations, R88C and R416W. We then evaluated mitochondrial function upon sequential injection of metabolic drugs. All three GFAP mutations caused significantly lowered basal respiration compared to GFAP WT cells (Figure 3A), confirming lower oxidative phosphorylation rates. GFAP mutations also resulted in a decreased maximal respiration upon addition of the uncoupler FCCP (Figure 3B), and a decrease in ATP production, represented by the decrease in Oxygen Consumption Rate (OCR) upon Oligomycin injection (Figure 3C).

**Figure 3.**
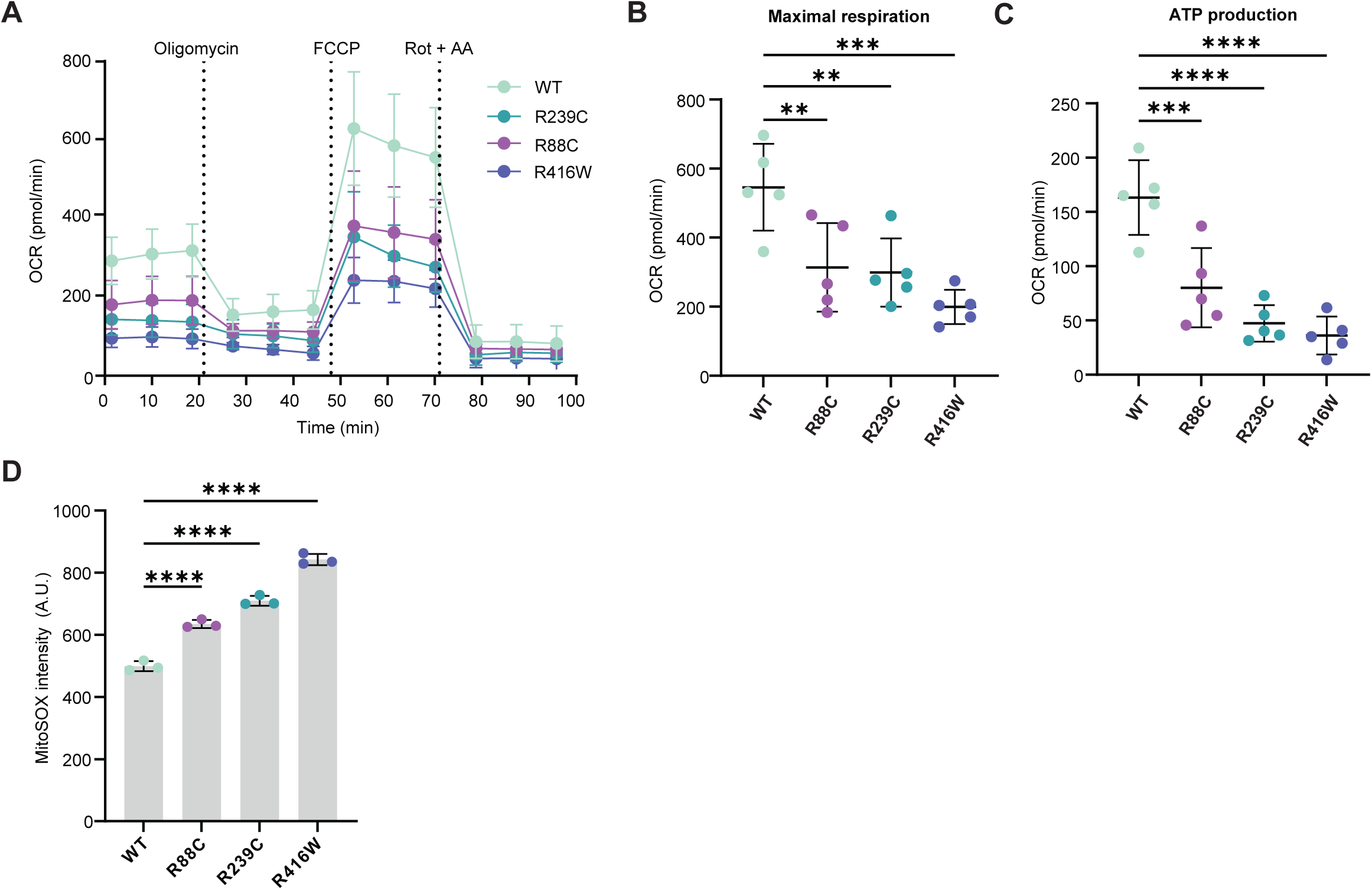
AxD GFAP mutations cause Electron Transport Chain dysfunction. **(A)** Seahorse mitochondrial stress test shows a decrease in oxygen consumption rate (OCR) in U87 GFAP R239C, R88C and R416W cells compared to WT. **(B)** GFAP mutations result in lower maximal respiration, calculated by the maximal OCR upon FCCP injection as well as a **(C)** decrease in ATP production measured by the decrease in OCR upon Oligomycin injection. **(D)** GFAP mutations result in increased superoxide production, measured by MitoSOX intensity. Significance determined by one-way ANOVA, with Benjamini Hochberg multiple testing correction comparing all mutant conditions to WT control. *** p < 0.001, **** p < 0.0001.

Superoxide is produced as a side-reaction at several sites of the ETC, including Complex I (Dlasková et al. 2008) and Complex III (Orr et al. 2015). Dysfunctional ETC assembly can result in increased superoxide production, as electrons may leak from the respiratory chain after which they can react with oxygen to form superoxide. Moreover, protons may leak back across the inner mitochondrial membrane (Brand et al. 2004). All tested GFAP mutations increased superoxide production compared to WT controls, as measured by the mitochondrial superoxide dye MitoSOX (Figure 3D). Together, these findings suggest that GFAP mutations induce compensatory upregulation of respiratory chain assembly machinery that is insufficient to restore efficient oxidative phosphorylation, ultimately resulting in electron transport chain dysfunction and mitochondrial oxidative stress.

### AxD organoids undergo a metabolic switch, fueling glycolysis

Glycolysis, the conversion of glucose to pyruvate and ultimately lactate, acts as an alternative metabolic pathway for ATP generation, while lactate produced by astrocytes serves as an important metabolic substrate for neurons. In the AxD organoids, an increase in most enzymes of glycolysis was observed, except for the neuron-specific ENO2 isoform, which could be explained by the lower number of neurons in AxD organoids compared to controls (Matusova et al. 2025). Similarly, an increase in pyruvate dehydrogenase (PDH) complex, and tricarboxylic acid (TCA) cycle enzymes were observed (Figure 4A). At the same time, all major components of the pentose phosphate pathway (PPP) were downregulated in AxD organoids. The PPP branches from the glycolysis pathway at the intermediate metabolite glucose-6-phosphate and contributes to NADPH and nucleotide production (TeSlaa et al. 2023). In contrast to the PPP, de novo serine biosynthesis, that uses the glycolysis intermediate metabolite 3-phospholglycerate, remains similar between WT and mutant cells. Together, these data suggest that in AxD glycolysis is directed towards feeding both the TCA cycle and subsequent oxidative phosphorylation.

**Figure 4.**
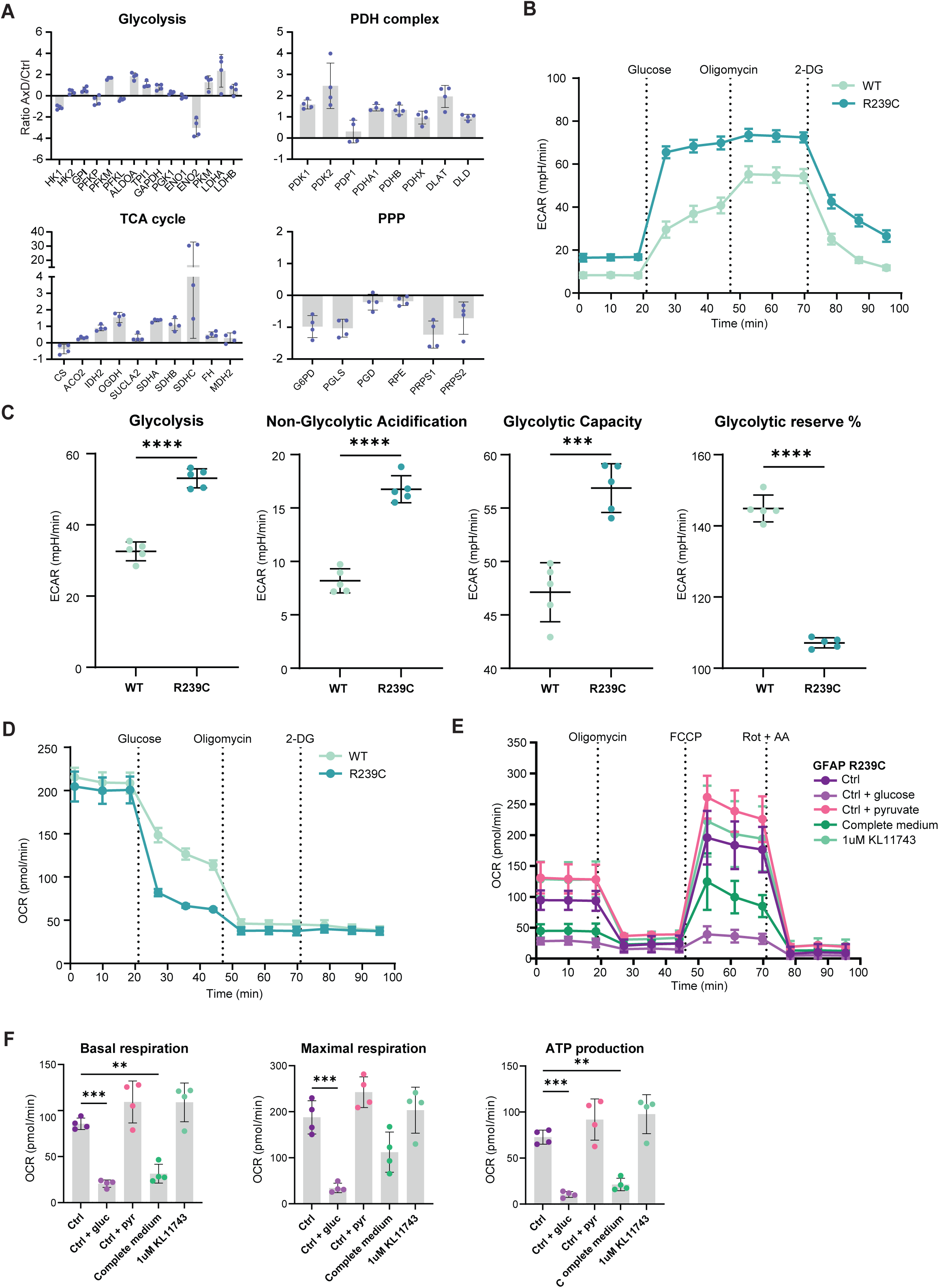
AxD organoids undergo a metabolic switch, fueling glycolysis. **(A)** AxD cortical organoids have increased abundance of enzymes involved in glycolysis, pyruvate dehydrogenase (PDH) complex, and the tricarboxylic acid (TCA) cycle, but decreased abundance of pentose phosphate pathway (PPP) enzymes. **(B)** Extracellular acidification rate profile of GFAP R239C and WT cells during the seahorse glycolysis stress test. **(C)** GFAP R239C cells show increased glycolysis and non-glycolytic acidification, and glycolytic capacity, while glycolytic reserve is decreased compared to WT, indicating that the GFAP R239C cells are operating at maximal capacity. **(D)** Oxygen consumption rate profile of GFAP R239C and WT cells during the seahorse glycolysis stress test. **(E)** Mito Stress Test OCR profile of GFAP R239C cells incubated with different media compositions and in combination with the glucose transporter inhibitor KL11743. **(F)** GFAP R239C shows decreased basal respiration, maximal respiration and ATP production in media containing glucose. ** p < 0.01, *** p < 0.001, **** p < 0.0001, determined by t-test (panel C) or one-way ANOVA with Benjamini-Hochberg correction for multiple testing.

As abundance does not necessarily reflect activity, we performed a Glycolysis Stress Test, where the Extracellular Acidification Rate (ECAR) is measured upon serial injections of glucose, oligomycin, and 2-Deoxy-D-glucose (2-DG). GFAP R239C cells showed both an increase in glycolysis, as well as non-glycolytic acidification, and had increased glycolytic capacity compared to WT controls (Figure 4B,C). Moreover, the mutant cells did not show an increase in ECAR upon oligomycin injection, indicating that they were already performing glycolysis at their maximal capacity. Basal OCR, measured in limited substrate media without glucose or pyruvate, but with L-glutamine, was not affected in GFAP R239C cells, but drastically decreased upon glucose injection (Figure 4D). We assessed mitochondrial function in different media conditions using the Mito Stress Test and found that glucose, and not pyruvate, is responsible for the decrease in OCR (Figure 4E). The addition of glucose caused a decrease in basal respiration, maximal respiration, and subsequent ATP production, which could be rescued by the addition of the glucose uptake inhibitor KL11743 (Figure 4F).

### GFAP R239C cells show increased glycerol-3-phosphate shuttle activity

To gain further insight into the metabolic adaptations caused by the GFAP R239C mutation, we performed MS-based metabolomics on the R239C mutant and WT cell lines (Figure 5A). Consistent with the metabolic flux analysis, GFAP R239C cells displayed increased abundance of multiple glycolytic and TCA cycle intermediates, confirming increased glycolysis and TCA cycle activity. As sustained glycolysis requires continuous regeneration of cytosolic NAD+, we specifically examined the two major NADH shuttle systems: the malate-aspartate shuttle (MAS) (Figure 5B) and glycerol-3-phosphate (G3P) shuttle (Figure 5C). Several MAS intermediates were increased in GFAP R239C cells, including malate, α-ketoglutarate (αKG), and glutamate (Figure 5A,S2). This pattern is consistent with enhanced cytosolic NADH oxidation through the MAS and is supported by the increased abundance of GOT1 observed in the proteomics dataset (Figure 5D). No difference in abundance was observed for the other MAS enzymes.

**Figure 5.**
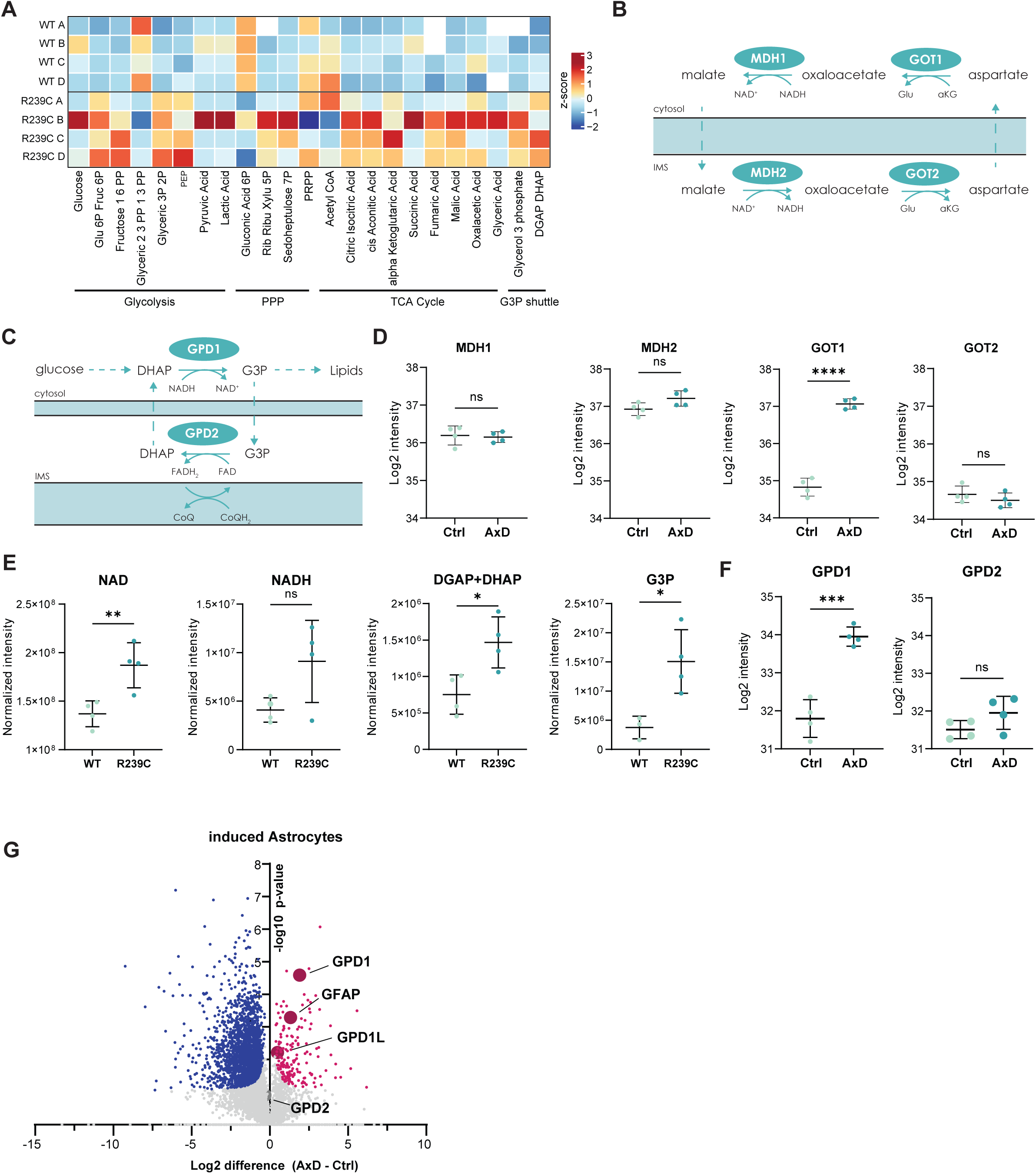
GFAP R239C cells show increased glycerol-3-phosphate shuttle activity. **(A)** MS-based analysis of polar metabolites comparing U87 GFAP R239C and WT cells. **(B)** Schematic of the Malate Aspartate Shuttle. **(C)** Relative abundance of malate aspartate shuttles enzymes in cortical AxD and isogenic control organoids. **(D)** Schematic of the Glycerol-3-Phosphate shuttle. **(E)** Intensity of G3P shuttle metabolites in GFAP WT and R239C U87 astrocytoma cells. **(F)** Relative abundance of G3P shuttle enzymes in cortical AxD and isogenic control organoids. **(G)** Vulcano plot comparing the proteome of induced astrocytes derived from patient-derived iPSCs and isogenic control line. Significantly upregulated proteins are highlighted in pink, downregulated proteins in blue.

Changes associated with the G3P shuttle were even more pronounced. Metabolites downstream of glyceric-2,3-phosphate/1,3-phosphate (GAPDH) were reduced, whereas both dihydroxyacetone phosphate (DHAP) and glycerol-3-phosphate (G3P) accumulated (Figure 5A,E). In the cytosolic arm of the G3P shuttle, GPD1 reduces DHAP to G3P while regenerating NAD+, thereby supporting continued glycolytic flux. G3P can subsequently be oxidized by mitochondrial GPD2, generating FADH2 and transferring electrons to the ETC through complex II (Figure 5D). Consistent with increased utilization of this pathway, GFAP R239C cells showed elevated fumarate and malate levels without a corresponding increase in succinate (Figure 5A), suggesting enhanced flux through complex II-associated metabolism. Furthermore, bulk proteomics data of AxD cortical organoids revealed a significant increase in GPD1 abundance, whereas GPD2 levels remained unchanged compared to isogenic control organoids (Figure 5F). To determine whether this alteration was astrocyte-specific, we performed bulk proteomics on induced astrocytes derived from the same patient-derived AxD iPSC line and its CRISPR-corrected isogenic control. Similar to the organoids, AxD astrocytes showed increased abundance of both GPD1 and the closely related GPD1L protein, whereas GPD2 levels were unaffected (Figure 5G).

Together, these metabolomic and proteomic findings identify enhanced G3P shuttle activity as a metabolic adaptation in GFAP R239C cells. The preferential upregulation of the cytosolic enzyme GPD1 relative to mitochondrial GPD2 suggests increased activity of the cytosolic part of the shuttle. Given that the G3P shuttle bypasses complex I and donates electrons directly to the ubiquinone pool via complex II, increased utilization of this pathway may represent a compensatory mechanism to maintain redox balance and cellular respiration in the context of ETC dysfunction.

### GFAP R239C cells partially uncouple the G3P shuttle to regenerate NAD+ for glycolysis

To investigate the role of the G3P shuttle in AxD in more detail, we treated our cellines with siRNA for GPD1 and assessed mitochondrial function (Figure S4A). Knockdown of GPD1 decreased oxygen consumption, spare respiratory capacity, maximal respiration and ATP production in GFAP R239C cells (Figure 6A,B), but not in WT (Figure S4B), indicating an essential role of the G3P shuttle in ETC function in GFAP R239C cells. Basal ECAR was increased upon GPD1 knockdown (Figure 6C), suggesting a compensatory shift towards lactate fermentation to regenerate NAD+ when G3P production is impaired. In contrast, superoxide levels were unaffected by GPD1 knockdown (Figure 6D), indicating that increased oxidative stress in GFAP R239C cells is not directly dependent on G3P shuttle activity.

**Figure 6.**
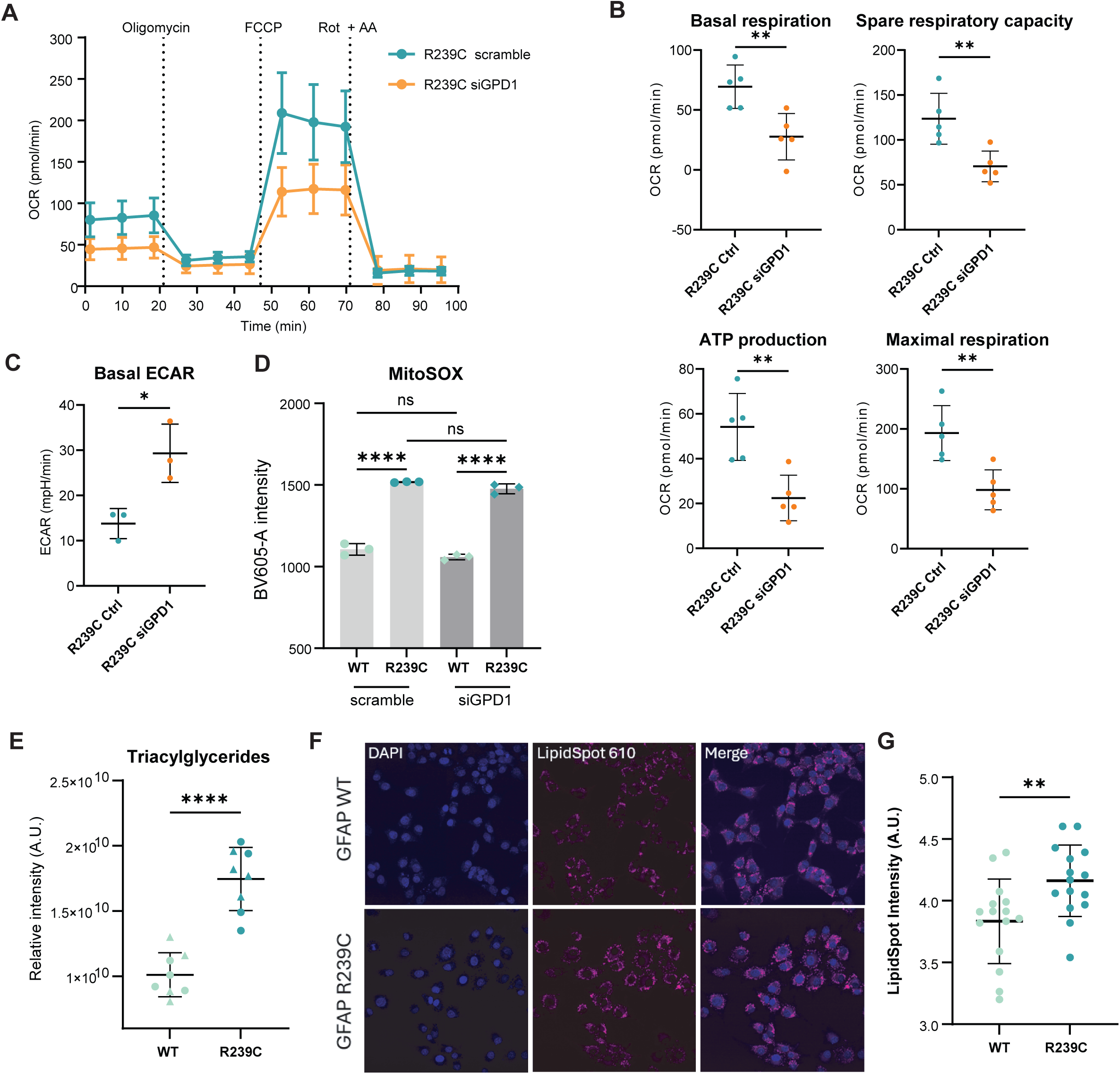
GFAP R239C cells partially uncouple the G3P shuttle to regenerate NAD+ for glycolysis. **(A)** Mito Stress Test OCR profile of GFAP R239C cells upon knockdown of GPD1. **(B)** Basal respiration, spare respiratory capacity, ATP production, and maximal respiration decrease significantly when GPD1 is knocked down. **(C)** Basal ECAR increases when GPD1 is knocked down in GFAP R239C cells. **(D)** Superoxide production in GFAP WT and R239C cells upon knockdown of GPD1. **(E)** MS-based lipidomics quantification of triacylglycerides in U87 GFAP WT and GFAP R239C astrocytoma cell lines. Different shapes indicate different clonal cell lines. **(F)** Visualization of lipid droplets using LipidSpot610. **(G)** Quantification of lipid droplets in GFAP WT and R239C cells.

Recent studies have highlighted an additional role of the G3P pathway beyond the canonical transfer of reducing equivalents to the electron transport chain. In kidney cancer cells, ETC dysfunction was shown to cause partial uncoupling of the G3P shuttle, with cytosolic GPD1 activity outpacing mitochondrial GPD2. This enables sustained regeneration of cytosolic NAD+ to support glycolysis while simultaneously diverting G3P towards lipid synthesis (Yao et al. 2023). Similarly, G3P biosynthesis has been identified as an important mechanism for NAD+ regeneration under conditions of ETC inhibition and hypoxia (Liu et al. 2021). Given the reduced respiratory capacity and increased glycolytic dependence of GFAP R239C cells, we hypothesized a similar mechanism in AxD.

If the cytosolic arm of the G3P shuttle becomes relatively more active than its mitochondrial counterpart, accumulation of G3P would be expected. Because G3P serves as the glycerol backbone for glycerolipid synthesis, excess G3P should promote triacylglyceride (TAG) production and lipid storage. To test this hypothesis, we performed MS-based lipidomics on GFAP R239C and WT cells. Indeed, GFAP R239C cells contained significantly higher levels of TAGs (Figure 6E). Moreover, at least part of these lipids were stored in lipid droplets, as demonstrated by increased LipidSpot intensity (Figure 6F,G). Together, these findings support a model in which ETC dysfunction promotes partial uncoupling of the G3P shuttle in GFAP R239C cells, allowing enhanced cytosolic NAD+ regeneration to sustain glycolytic flux while diverting excess G3P towards lipid synthesis and storage.

## Discussion

A major challenge in understanding AxD pathogenesis is distinguishing mutation-specific gain-of-function effects from the consequences of progressive GFAP accumulation. Mutant GFAP not only perturbs cellular homeostasis, but also appears to promote GFAP accumulation through a self-reinforcing stress response (Escartin et al. 2021; Heaven et al. 2019). As a result, disease-associated phenotypes may arise from the direct effects of the mutation, from increased GFAP abundance itself, or from downstream compensatory adaptations. Disentangling these mechanisms has proven difficult using conventional bulk approaches but is critical for understanding the molecular basis of AxD.

Here, we used astrocytoma cell lines overexpressing either WT GFAP or a common AxD-causing GFAP R239C mutant, both fused to a GFP reporter. Rather than comparing cell populations as bulk samples, we exploited variation in GFP intensity between individual cells as a proxy for GFAP abundance using single cell proteomics. We isolated single cells spanning a broad range of GFAP expression levels and used MS-based single cell proteomics to analyze their individual proteomes, which allowed us to investigate and correlate proteomic responses across a range of GFAP abundances and to distinguish alterations associated with increasing GFAP levels from those specifically induced by the R239C mutation. Such discrimination is difficult to achieve using conventional bulk proteomic approaches, where the effects of protein abundance and mutation status are inherently confounded. By resolving cellular heterogeneity at the single-cell level, we were therefore able to separate dosage-dependent effects from mutation-specific responses, providing a more refined understanding of AxD pathophysiology.

Indeed, cells expressing high levels of WT GFAP increasingly resembled R239C-expressing cells at the proteomic level. These observations provide support for the toxic-threshold model of AxD, which proposes that pathology emerges when GFAP accumulates beyond a critical level. At the same time, our single-cell approach reveals that this model alone cannot fully explain disease pathogenesis. While several phenotypes previously attributed to mutant GFAP seem largely driven by GFAP abundance, other alterations remain specifically associated with the R239C mutation. Increased abundance of mitochondrial antioxidant enzymes was highly correlated with mutant but not WT GFAP abundance. Similarly, mutant GFAP levels correlated with 11S and immunoproteasome subunit abundance, which was confirmed by bulk proteomics data of patient-derived AxD cortical organoids. Together, these results point towards the remodeling of the proteasome to facilitate the removal of oxidized proteins in an ATP and ubiquitin-independent manner. Mutant GFAP has been shown to inhibit proteasomal function, and 20S subunits have been found trapped in Rosenthal Fibers (Tang et al. 2010). Moreover, an increase in ubiquitinated GFAP and other proteins has been reported in several models of AxD (Lin et al. 2024; Lin and Perng 2026; Yang et al. 2022). Our data suggests an additional mechanism, in which at least part of the proteasome pool is remodeled towards the degradation of potentially harmful oxidized proteins, thereby prioritizing ubiquitin-independent degradation. This might explain the buildup of ubiquitinated proteins in AxD models but needs further investigation.

Our data also indicate mitochondrial dysfunction as a consequence of mutant GFAP expression. Our SCP approach revealed an increased correlation between mutant GFAP and subunits of complex V of the ETC. We previously performed bulk proteomics analyses on 150-day old AxD cortical organoids. These data showed a vast increase in nuclear-encoded mitochondrial proteins in AxD organoids compared to isogenic control organoids, without a corresponding increase in mitochondrially encoded proteins. The proteins showing the strongest changes were not primarily catalytic ETC subunits, but rather assembly factors involved in respiratory complex biogenesis. Because respiratory complexes require the coordinated incorporation of subunits encoded by both the nuclear and mitochondrial genomes, this pattern may indicate impaired coordination of ETC assembly and maintenance. This would be consistent with a compensatory cellular response attempting to maintain respiratory chain function despite incomplete or unstable complex formation. In contrast to our data, a recent study on rats carrying the R237H mutation reported reductions in mitochondrial DNA-encoded proteins of at day 56, but not yet at day 21 (Berman et al. 2025). Thus, temporal progression may account for some discrepancies in observations. Moreover, important differences between species exist in GFAP biology, as human GFAP expression begins during corticogenesis whereas expression in rodents occurs largely after corticogenesis has been completed (Dykstra et al., 2025). Finally, regional differences in the analyzed tissues may further contribute to divergent observations across studies.

Additional evidence for ETC dysfunction comes from the selective enrichment of complex II components. Unlike complex I, complex II transfers electrons from FADH2 directly into the ubiquinone pool and therefore provides a mechanism to support respiratory chain activity while partially bypassing complex I-dependent electron transfer. Increased reliance on complex II may therefore represent an adaptive response that partially bypasses deficiencies elsewhere in the respiratory chain. At the same time, we observed increased abundance of proteins associated with both the MAS and G3P shuttles, indicating enhanced efforts to transfer cytosolic reducing equivalents into mitochondria. Together, these findings suggest that cells continue to supply substrates to the ETC despite compromised respiratory efficiency.

The notion that ETC dysfunction is an upstream event is further supported by our metabolic flux measurements. Although oxygen consumption rates were reduced in AxD cells, these differences disappeared in glucose-free medium, suggesting that the observed respiratory phenotype is not caused by an absolute inability to consume oxygen but rather by a cellular shift away from oxidative metabolism. Simultaneously, we observed an increase in glycolysis and enhanced glycolytic capacity, together with a lower glycolytic reserve, indicating that the GFAP R239C cells are operating at full metabolic capacity. Such metabolic reprogramming is characteristic of cells experiencing limitations in oxidative phosphorylation. By increasing glycolytic flux, cells can maintain ATP production despite reduced mitochondrial efficiency.

Mitochondrial dysfunction has emerged as a recurring feature of AxD, with previous studies reporting increased oxidative stress and superoxide production, reduced oxygen consumption and ATP generation (Bellitto et al. 2025), altered mitochondrial morphology (Hernandez-Gerez et al. 2026; Xiong et al. 2025), impaired mitochondrial transfer between astrocytes and neurons (Gao et al. 2019), and evidence for a shift away from fatty acid oxidation (Bellitto et al. 2025). Our findings support this growing body of literature and suggest that dysfunction of the electron transport chain may, at least partially, explain these metabolic abnormalities.

Although GFAP R239C cells displayed increased glycolytic activity, their metabolic phenotype differs from a classical Warburg response. In the Warburg effect, pyruvate is converted to lactate to regenerate cytosolic NAD⁺ and sustain glycolytic flux, but at the expense of mitochondrial oxidation of pyruvate and TCA cycle activity (Luengo et al. 2021). In contrast, GFAP R239C cells showed only a limited increase in lactate production while displaying increased TCA cycle intermediates, increased abundance of TCA cycle enzymes, and the partially uncoupled G3P shuttle. This may be particularly important in astrocytes, where lactate functions not only as a glycolytic product, but also as a key metabolic substrate for neurons through the astrocyte-neuron lactate shuttle (Magistretti and Allaman 2018). By regenerating NAD^+^ through the G3P shuttle, astrocytes may preserve lactate availability for neuronal support while maintaining glycolytic flux. Moreover, although oxygen consumption was reduced under standard culture conditions, this difference disappeared in glucose-free medium, indicating that the lower OCR is not caused by an absolute inability of the mitochondria to consume oxygen. Instead, GFAP R239C cells appear to preferentially utilize glycolysis when glucose is available while maintaining active mitochondrial carbon metabolism. These observations suggest that NAD⁺ regeneration occurs primarily through mitochondrial shuttle systems rather than lactate fermentation, allowing glycolysis and TCA cycle activity to operate in parallel despite evidence of electron transport chain dysfunction.

Sustained glycolysis requires continuous regeneration of cytosolic NAD+, and our data suggest that in AxD activation of the G3P shuttle may play an important role in this process. Compared to the MAS, the G3P shuttle remains relatively understudied, although its importance has gained traction, for instance as a backup system securing metabolic flexibility in neurons (Dhoundiyal et al. 2022). In the canonical shuttle, cytosolic GPD1 reduces DHAP to G3P, thereby regenerating NAD⁺ required for glycolysis. G3P is subsequently oxidized back to DHAP by the mitochondrial enzyme GPD2, transferring electrons to FAD and ultimately to the ETC. Increased abundance of proteins associated with both the MAS and G3P shuttles suggests enhanced efforts to maintain redox balance under conditions of impaired oxidative phosphorylation. However, activation of the G3P pathway was accompanied by marked accumulation of triacylglycerides and lipid droplets. Because G3P serves not only as a shuttle intermediate but also as the backbone for glycerolipid synthesis, its accumulation suggests that a fraction of the metabolic flux is redirected away from mitochondrial oxidation. Recent studies demonstrated that ETC dysfunction can promote partial uncoupling of the G3P shuttle, favoring cytosolic GPD1 activity to regenerate NAD+ and sustain glycolysis while reducing the transfer of reducing equivalents into mitochondria and diverting G3P toward lipid synthesis (Yao et al. 2023). Furthermore, G3P production has been shown to support NAD+ regeneration during hypoxia and ETC inhibition (Liu et al. 2021). Our findings are highly consistent with such a mechanism. The concurrent increase in glycolysis, triacylglycerides, and lipid droplets observed here is therefore consistent with a model in which ETC dysfunction drives partial uncoupling of the G3P shuttle. In this scenario, enhanced GPD1 activity maintains glycolytic flux through NAD⁺ regeneration, while excess G3P is increasingly utilized for lipid synthesis and storage rather than mitochondrial electron transfer.

Our model therefore provides a mechanistic link between ETC dysfunction, enhanced glycolysis, and the previously reported lipid metabolic alterations in AxD. Rather than representing independent phenotypes, lipid droplet accumulation, increased triglyceride synthesis, reduced oxygen consumption, and enhanced glycolysis may arise from a common adaptation aimed at maintaining cytosolic NAD⁺ levels and ATP production when mitochondrial electron transport becomes limiting. Increased synthesis of triacylglycerides and free fatty acids in an environment of elevated oxidative stress would furthermore be expected to promote previously reported lipid peroxidation (Viedma-Poyatos et al. 2022).

An important unresolved question is how mutant GFAP leads to ETC dysfunction. Recent evidence suggests that GFAP participates in Drp1-mediated mitochondrial fission, and that the disease-associated GFAP R88C and R416W mutations resulted in fragmented mitochondria (Xiong et al. 2025). Together with reports of elongated mitochondria in R239C models (Viedma-Poyatos et al. 2022), these findings raise the possibility that abnormalities in mitochondrial remodeling impair the assembly, maintenance, or turnover of respiratory complexes. Although this hypothesis remains speculative, it provides a plausible mechanistic link between mutant GFAP accumulation and the respiratory chain abnormalities observed.

In conclusion, our study demonstrates the power of single-cell proteomics to resolve the complex relationship between GFAP abundance and mutation-specific pathology in AxD. By separating these effects, we identify ETC dysfunction as a potential player linking GFAP mutation to oxidative stress, proteasome remodeling, metabolic rewiring and lipid accumulation. We further uncover the G3P pathway as a component of the astrocytic response to GFAP-associated stress. These findings not only advance our understanding of AxD pathogenesis but also provide new insight into the mechanisms by which astrocyte dysfunction can shape neural vulnerability and disease progression more generally.

## Author’s contributions

CAGHVG, EMH and BMTB conceived the study, BMTB and EMH guided and supervised the project, CG and ECAS performed and analyzed the lipidomics and metabolomics experiments, CAGHVG, ED, SEB and YA performed all other experiments, CAGHVG, PSA, RMVE and HRV analyzed the proteomics data and prepared the related figures. CAGHVG and BMTB prepared the manuscript and figures, all authors read and approved the final manuscript.

## Acknowledgements

We would like to thank Maaike van Heeswijk and Fleur Ackermann with their help with cloning, Huanjie Huang and Livio Kleij with help with live imaging experiments, and all CMM lab members for valuable discussions and feedback.

---

**Figure S1.**
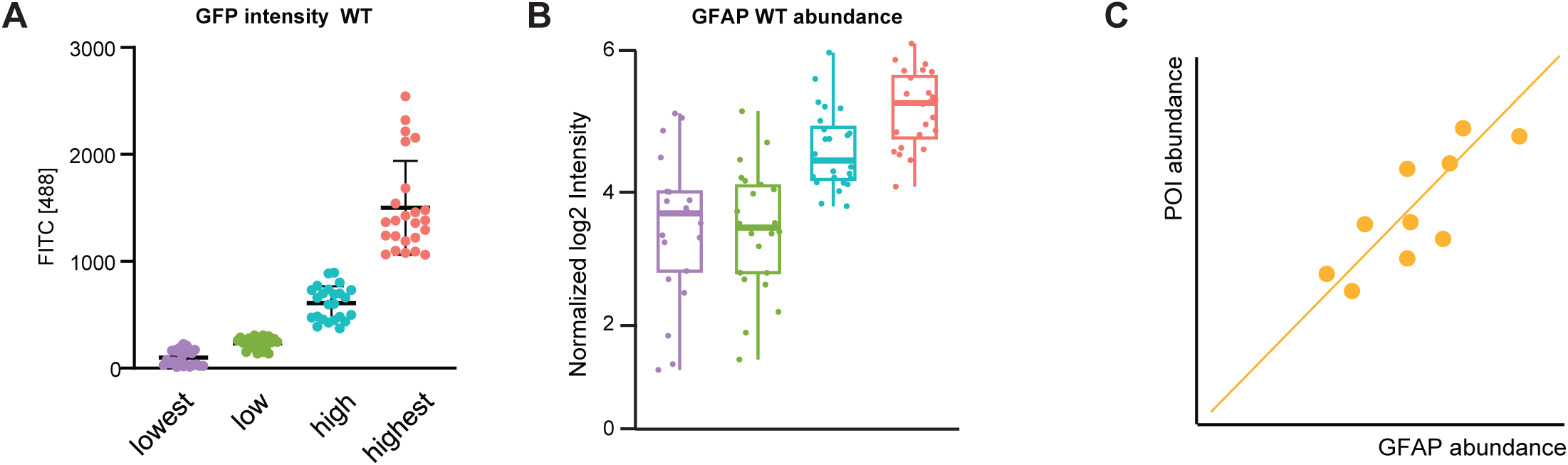
With figure 1. **(A)** Single cells were sorted based on eGFP-GFAP intensity into four bins (n=24 per bin). **(B)** Mass spectrometry-based quantification of GFAP protein of the single cell samples. **(C)** To distinguish proteomic alterations caused by increased GFAP abundance from perturbations caused by the GFAP R239C mutation, we calculated the correlation between the abundance of GFAP and all other identified proteins in single cells. Each dot represents a single cell.

**Figure S2.**
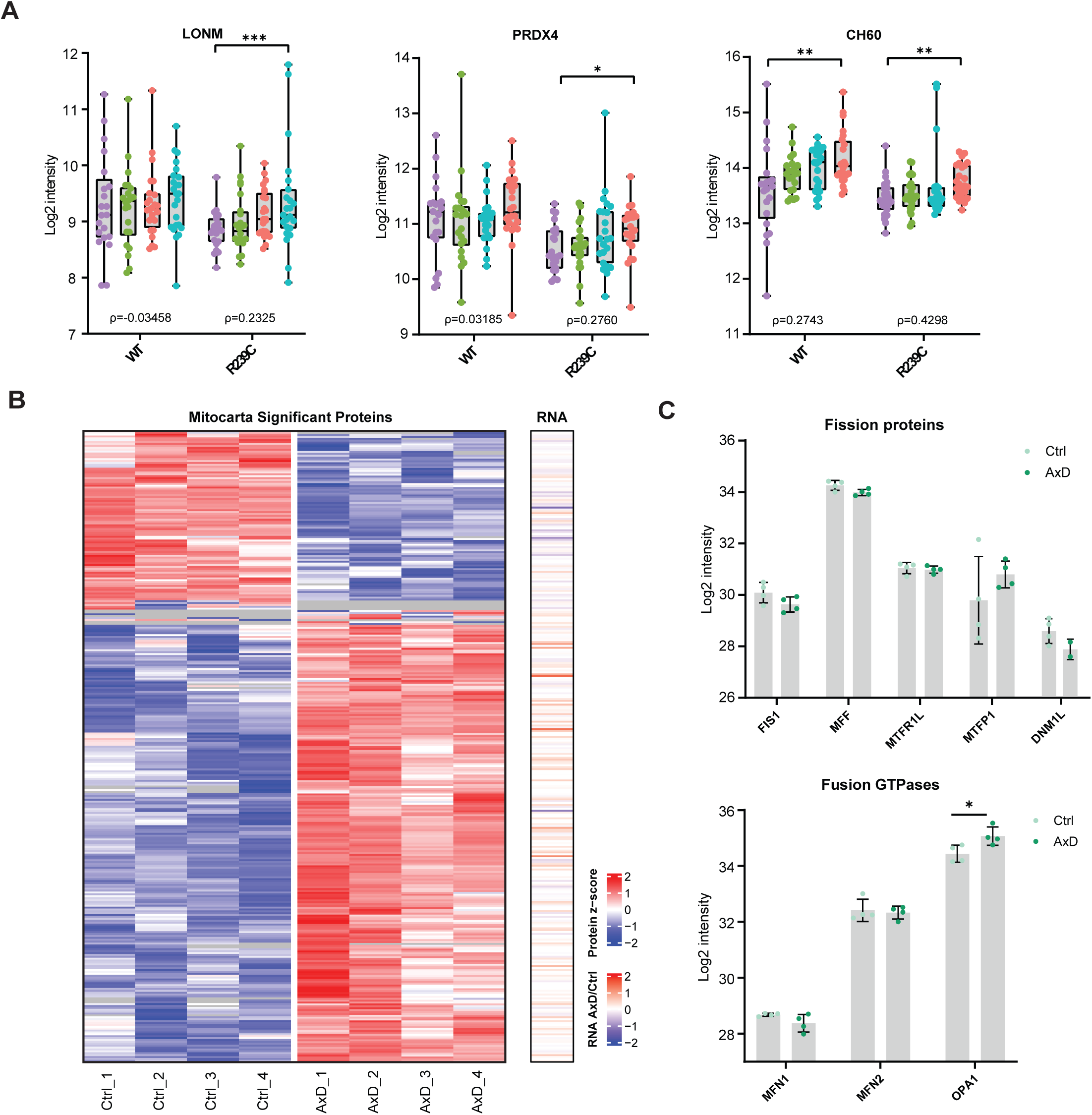
With figure 2. **(A)** Single cell protein abundances of proteins involved in mitochondrial redox balance in U251 GFAP WT and R239C cells. **(B)** Heatmap of all mitochondrial proteins with significantly altered abundance in AxD cortical organoids compared to isogenic controls. The additional RNA column indicates transcript ratio in AxD versus control from pseudobulk transcriptomics data of 165-day old cortical organoids data (Matusova et al. 2025). **(C)** Relative abundance of proteins involved in mitochondrial fission and fusion detected in whole proteome analysis of AxD and isogenic control cortical organoids.

**Figure S3.**
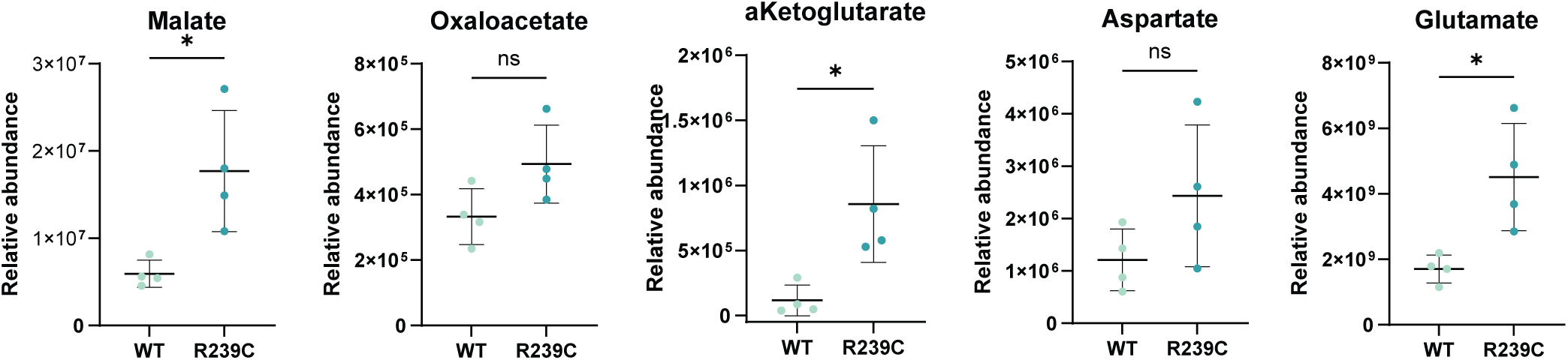
With figure 5. Relative abundance of polar metabolites involved in the malate aspartate shuttle. * p < 0.05, determined by t-test.

**Figure S4.**
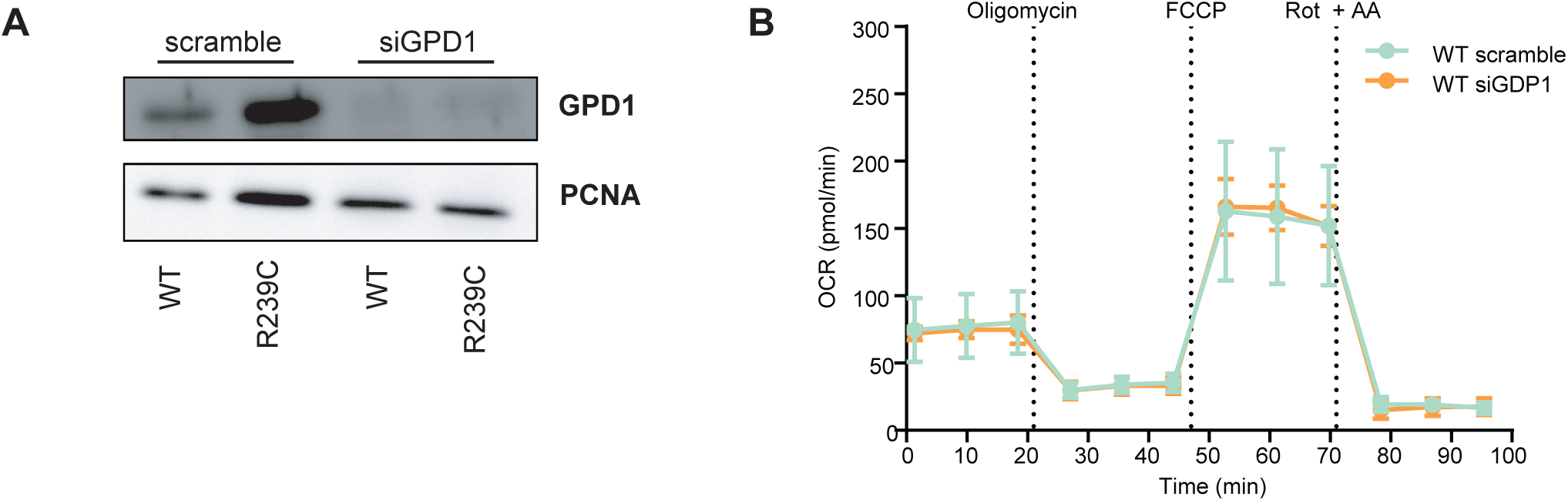
With figure 6. **(A)** Western blot analysis of GPD1 upon siGPD1 treatment. PCNA was used as a loading control. **(B)** Mito Stress Test OCR profile of GFAP WT cells upon knockdown of GPD1.

